# A comprehensive HA-tagged PRC1 toolkit enables standardized chromatin profiling and proteomic analysis in *Drosophila melanogaster*

**DOI:** 10.64898/2026.07.27.741006

**Authors:** Lauriane Fritsch, Vincent Loubiere, Axelle Donjon, Julia Morales-Sanfrutos, Eduard Sabidó, Anne-Marie Martinez, Bernd Schuettengruber, Giacomo Cavalli

## Abstract

Polycomb group (PcG) proteins are evolutionarily conserved epigenetic regulators that maintain transcriptional states during development and are frequently misregulated in disease. Here, we generated a comprehensive collection of endogenously HA-tagged alleles for the core components of Polycomb Repressive Complex 1 (PRC1) and additional chromatin regulators in *Drosophila melanogaster* using CRISPR/Cas9-mediated genome engineering. We show that endogenous HA tagging preserves protein expression, chromatin localization, and genome-wide binding profiles, enabling direct comparison using a common antibody to analyze distinct PcG subunits. Proteomic analyses recovered known components of canonical and non-canonical PRC1 and PRC2 complexes and identified additional PcG-associated factors. Notably, multiple components of the nuclear pore complex (NPC) were detected as PcG interactors, and genetic analyses confirmed a functional interaction between them. This resource provides a platform for standardized analysis of Polycomb function and chromatin regulation that can easily be extended to other chromatin regulatory complexes and applied across different developmental stages.

**Article Summary:** Polycomb group (PcG) proteins are essential regulators of development and genome function. Comparative studies of individual PcG components are often limited by the availability and quality of specific antibodies. Here, we generated a collection of *Drosophila melanogaster* lines carrying endogenous HA epitope tags in core components of the canonical Polycomb Repressive Complex 1 (PRC1), along with additional chromatin regulators. These tagged proteins faithfully reproduce native chromatin binding and can be used for chromatin mapping and protein interaction studies using a single, highly specific antibody. This standardized resource provides a practical tool for investigating PcG function and chromatin regulation in vivo.

## Introduction

Epigenetic mechanisms, primarily mediated through histone and DNA modifications, play a pivotal role in orchestrating cell fates and their response to environmental cues (CAVALLI AND HEARD 2019). One of the most important epigenetic regulators comprises the evolutionary conserved Polycomb group (PcG) proteins, originally identified in *Drosophila melanogaster* as an epigenetic memory system that maintains the repressed state of homeotic (HOX) genes during *Drosophila* development (LEWIS 1978). After decades of research, PcG proteins are now well recognized as key factors of genome regulation with a pleiotropic role in the regulation of genes involved in development, growth control, major signaling pathways, differentiation, cell-fate determination and tumorigenesis (SCHUETTENGRUBER *et al*. 2017; CHAN AND MOREY 2019; PARRENO *et al*. 2022; MARTINEZ 2025). In *Drosophila*, PcG proteins act as large multimeric complexes. The two main complexes are the Polycomb repressive complex 2 (PRC2) and the Polycomb repressive complex 1 (PRC1) (reviewed in (SCHUETTENGRUBER *et al*. 2017)). PRC2 contains the core subunits Enhancer of zeste (E(Z)), Suppressor of zeste 12 (SU(Z)12), Extra sex combs (ESC), and NURF55/CAF1. E(Z) has a methyltransferase activity mediating the methylation of the lysine 27 of histone H3 (H3K27me3), which is a hallmark of PcG-mediated gene silencing (CZERMIN *et al*. 2002; MULLER *et al*. 2002). PRC1 contains the core subunits Polycomb (PC), Polyhomeotic (PH), Sex combs extra (SCE, also known as dRING), and Posterior sex combs (PSC) subunits. dRING is an E3 ubiquitin-protein ligase that catalyzes the monoubiquitination of the lysine 118 of histone H2A (H2AK118ub), contributing to PRC2 recruitment, chromatin compaction and gene silencing (SHAO *et al*. 1999; FRANCIS *et al*. 2001). Furthermore, non-canonical Polycomb complexes exist for both PRC1 and PRC2 in mammals and in *Drosophila* (KING *et al*. 2005; KASSIS *et al*. 2017). In flies, canonical PRC1 (cPRC1) and non-canonical PRC1 (ncPRC1) share the E3 ubiquitin ligase dRING but associate with different PCGF family proteins: the cPRC1 contains Posterior sex combs (PSC) or Suppressor of zeste 2 (SU(Z)2), while ncPRC1 contains Lethal (3) 73Ah (L(3)73Ah) (KING *et al*. 2005; KANG *et al*. 2022). Affinity purification of L(3)73Ah from formaldehyde cross-linked *Drosophila* embryo nuclei identified additional components orthologous to subunits of mammalian PRC1.1 and PRC1.3/5 complexes, including RYBP, which is a shared component of all ncPRC1 assemblies (KANG *et al*. 2022).

*Drosophila* PRC2 also exists in multiple forms through association with distinct accessory proteins: PRC2.1 contains Polycomb-like (PCL), whereas PRC2.2 incorporates JARID2 and AEBP2 (also known as JING) into the core complex. These accessory subunits modulate PRC2 chromatin binding and/or catalytic activity, thereby contributing to functional diversification of PRC2 during development (NEKRASOV *et al*. 2007; HERZ *et al*. 2012; KALB *et al*. 2014).

PcG complexes are recruited to their target sites by specific DNA sequences, called Polycomb response elements (PREs), via interactions with sequence-specific DNA-binding proteins (SIMON *et al*. 1993; CHAN *et al*. 1994; KASSIS *et al*. 2017).

Initial studies suggested a hierarchical recruitment model in which PRC1 recruitment depends on the presence of PRC2-mediated H3K27me3 (FISCHLE *et al*. 2003; WANG *et al*. 2004). This model was supported by the extensive colocalization of PRC1 and PRC2 complexes observed in *Drosophila* embryos (SCHWARTZ *et al*. 2006; SCHUETTENGRUBER *et al*. 2009).

However, PRC1 can bind to PREs even when H3K27me3 levels are strongly reduced or absent, indicating that PRC1 recruitment to PREs can occur independently of H3K27me3 in flies (LOUBIERE *et al*. 2016). Moreover, PRC1 contributes to stabilizing PRC2 occupancy and H3K27me3 domains, particularly at later developmental stages, supporting a cooperative rather than strictly hierarchical relationship between PRC1 and PRC2 (PAPP AND MULLER 2006; SCHUETTENGRUBER *et al*. 2014). PcG binding to PREs is also dynamically regulated during development, and many PcG binding sites are acquired during larval stages that are not occupied by PcG proteins earlier in development (LOUBIERE *et al*. 2016).

Most of our knowledge about PcG complex composition and their genomic localization during development stems from proteomic or genomic studies relying on antibodies directed against members of the core subunits and/or accessory proteins. Consequently, these analyses are restricted to proteins for which suitable antibodies are available. In addition, the results may be influenced by differences in antibody quality or specificity, while studies based on UAS/Gal4-mediated expression may introduce biases due to non-physiological expression levels. The advent of CRISPR/Cas9-based genome engineering has provided a powerful approach to easily and precisely manipulate the genome and modify or insert specific DNA sequences (BASSETT AND LIU 2014). For example, tagging different proteins of the same complex with an identical, small HA epitope allows standardized, side-by-side comparison of their chromatin occupancy, complex assembly and function, while minimizing confounding effects associated with antibody variability and non-physiological expression levels.

Here, we report the systematic tagging of endogenous genes coding for all members of the core cPRC1 complex, including different isoforms of PH using CRISPR/Cas9 genome engineering. In addition, we tagged SU(Z)12, a core component of the PRC2 complex, as well as other chromatin regulators that function independently of PcG proteins, providing a valuable toolkit to study the function of these chromatin regulators during *Drosophila* development. We show that tagging of the endogenous genes at their C-terminal end does not affect their expression levels, nor their function. Furthermore, the genomic localization of the tagged proteins is unaltered. To demonstrate the utility of this resource, we profiled HA-tagged members of cPRC1, ncPRC1, and PRC2 across different developmental stages using different mapping techniques, including chromatin immunoprecipitation followed by sequencing (ChIP–seq) and CUT&RUN. This enabled quantitative comparison of the genomic distribution of subunits from distinct complexes in *Drosophila* embryos or 3^rd^ instar imaginal discs and revealed that, during *Drosophila* development, cPRC1 and ncPRC1 largely occupy a common set of PREs. We further performed proteomic analyses using HA-tagged core members of PRC1 and PRC2, demonstrating that this approach not only enables the detection of all stoichiometric components of these PcG complexes but also the identification of additional chromatin regulators, such as components of the nuclear pore complex (NPC). Finally, we validated genetic interactions between PcG proteins and the NPC, confirming a functional interplay between these chromatin-associated complexes.

## Materials and Methods

### Fly strains and *Drosophila* genetics

Flies were raised on a standard cornmeal yeast extract medium at 25 °C unless otherwise indicated. Fly lines were obtained from the Bloomington *Drosophila* Stock Center (BDSC), and crosses were performed using the stocks indicated in **Supplementary Table 1**. Work involving transgenic *Drosophila* lines was performed under ethical approval no. n6906C2 of the Ministère de l’Enseignement Supérieur, de la Recherche et de l’Innovation, issued on 8 April 2020. The following knock-in lines were generated in this study: *Pc*^HA-Myc^, *Psc*^HA-Myc^, *Su(z)12*^HA-Myc^, *ph-p*^HA-Myc^, *ph-d*^HA-FLAG^, *Sce*^HA-FLAG^, and *Su(Hw)*^HA-FLAG^.

### gRNA selection and plasmid cloning

Target gene sequences were obtained from FlyBase and analyzed to find two optimal target gRNA sites before and after the stop codon. Selected gRNA sequences displayed no predicted off-target sites and specificity scores close to 100 (https://crispor.gi.ucsc.edu). Optimal target sites were then confirmed by sequencing the genomic loci from DNA extracted from the fly lines used for plasmid injection. Cloning gRNA expression vectors was done as previously described in (OGIYAMA *et al*. 2018).To generate C-terminal tagging HA-Myc we used the pHD-DsRed vector (addgene #51424) as template for homology-directed repair. C-terminal homology arms of the gene of interest were obtained by PCR and cloned into pHD-DsRed. N-terminal homology arms carrying the Ha-Myc tag sequence were obtained by gene synthesis and inserted into the pHD-DsRed donor plasmid (see **Supplementary Table 1**).

### Embryo injections

Embryo injections were performed according to standard procedures. For each construct, 300-500 embryos were injected into *y*^1^ *M{RFP[3xP3.PB] GFP[E.3xP3]=vas-Cas9}ZH-2A w*^1118^/*FM7c, w*^1118^; *PBac{y[+mDint2] GFP[E.3xP3]=vas-Cas9}VK00027* or *y w; nos-Cas9* (*attP40*) embryos (for CRISPR/Cas9 constructs). Surviving G0 adults were backcrossed individually to *w1118* flies, and transformants were screened for DsRed expression. Stable transgenic lines were subsequently established by crossing transgenic flies to the appropriate balancer stock. In a second round of selection, balanced progeny that had lost the chromosome carrying the Cas9 transgene were back-crossed to the appropriate balancer stock to establish stable lines. The embryo injections, and selection of DsRed-positive transformants for tagging *Sce*, *Su(Hw)*, and *ph-d* were performed by BestGene Inc (https://www.thebestgene.com). The design, injections and selection of DsRed-positive flies for tagging *Sce*, *Su(Hw)* and *ph-p* was performed by Rainbow Transgenic Flies Inc (https://www.rainbowgene.com/). The DsRed selection cassette was then excised by crossing the tagged lines to *y*^1^ *w*^67c23^ *P{y[+mDint2]=Crey}1b*; *sna*^Sco^/*CyO*; *Dr*^1^/*TM3, Sb*^1^ (BDSC #34516). For X-linked loci, the DsRed cassette was removed by crossing to *hs-cre CyO/sna Sco* (gift from Judith Kassis).

### Genotyping *Drosophila* strains

Correct insertions and genotypes were confirmed by PCR. Briefly, total DNA was extracted from 5 flies using puregene kit (Qiagen#158063). Genotyping PCRs were performed using primers annealing to the 5’ and 3’ ends of the flanking regions of the inserted tag (**Supplementary Table 1**).

### Genetic analysis of extra sex comb (*esc*) mutant phenotypes

Flies were reared and crossed at 21 °C. A total of 40 virgin *Pc*^1^ mutant females were crossed with 15 males of each genotype to be tested. Flies were allowed to lay eggs at 21 °C for 16 h, and the presence of extra sex combs on the T2 and T3 legs was quantified in heterozygous or double-heterozygous F1 flies. The precise genotypes of the flies used for the different analyses are listed below:

*Pc*^1^/*TM1* (BDSC #1728); *Nup98-96*^339^ (BDSC #4651)/ *Nup98-96*^G2120^ (BDSC #28433)/*Mtor*^k03905^ (BDSC #10537)/ *BEAF-32*^KG06304^ (BDSC #14058).

### Immunostaining experiments of polytene chromosomes

Salivary glands from wandering 3^rd^-instar larvae were dissected at room temperature (RT) in 1× PBS and processed as described in (W. SULLIVAN 2000). The following primary antibodies were used: rat anti-HA (1:200, Roche, catalogue no 11867423001) and rabbit anti-PC (SCHUETTENGRUBER *et al*. 2009). The following secondary antibodies were used: donkey anti-rat Alexa Fluor 488 (1:500, Invitrogen, catalogue no. A-21208) and donkey anti-rabbit 555 (1:500, Invitrogen, catalogue no. A-31572). Tissues were washed three times in PBTr. DAPI (4,6-diamidino-2-phenylindole) staining was performed at a final concentration of 1 µg ml−1 for 15 min. Images were acquired using an Apotome Z3 microscope equipped with a 60x oil-immersion objective. Images were processed using Fiji (ImageJ).

### CUT&RUN experiments

CUT&RUN experiments were performed as described by Kami Ahmad in protocols.io (AHMAD) with minor modifications. We dissected 20 eye-antennal imaginal discs (EDs) for histone mark profiling and 40 EDs for chromatin-bound factor profiling in Schneider’s medium, centrifuged the tissues for 3 min at 2000g and washed them twice with Wash+ buffer before adding Concanavalin A-coated beads. For experiments involving non-histone proteins, EDs were crosslinked with 0.1% formaldehyde for 5 min at RT on a rotating wheel. MNase digestion was performed using pAG-MNase Enzyme (Cell Signaling Technology) for 30 min at 4°C. Following reverse crosslinking for 4 h at 65°C and Proteinase K digestion, DNA was purified using SPRIselect beads and eluted in 50 μl of Tris-EDTA. DNA libraries for sequencing were prepared using the NEBNext Ultra II DNA Library Prep Kit for Illumina. Paired-end sequencing (150 bp; approximatively 2 Gb per sample) was performed by Novogene (https://en.novogene.com/). The following antibodies were used: anti-H3K27me3 (1:100, Active Motif, catalogue no. 39155), anti-H2AK118ub (1:100, Cell Signaling Technology, catalogue no. 8240), anti-HA (1:50, Cell Signaling Technology, catalogue no. 3724) and anti-PC (SCHUETTENGRUBER *et al*. 2009). All experiments were performed in biological duplicates.

### ChIP-seq experiments

ChIP–seq on embryos were performed as previously described (SCHUETTENGRUBER *et al*. 2009). When necessary, several batches of embryos collected over 16-20 h egg laying periods were pooled, frozen in liquid nitrogen, and stored at −80 °C until sufficient material was obtained. Chromatin was sonicated using a Covaris instrument (Diagenode) for 10 min in 130 µl tubes (peak power = 105W; duty factor = 10%; cycles per burst = 200). Antibodies against HA, PC (SCHUETTENGRUBER *et al*. 2009), SU(HW) (MOSHKOVICH AND LEI 2010) and SU(Z)12 (LOUBIERE *et al*. 2017) were used for immunoprecipitation at a dilution of 1:100. Following decrosslinking, DNA was purified using MicroChIP DiaPure columns (Diagenode). DNA libraries for sequencing were prepared using the NEBNext Ultra II DNA Library Prep Kit for Illumina. Paired-end sequencing (sequencing (150 bp; approximately 2 Gb per sample) was performed by Novogene (https://en.novogene.com/). All experiments were performed in biological duplicates.

### Processing and analysis of ChIP-seq and CUT&RUN sequencing data

Bioinformatic analyses were performed in R v4.5.0 (https://www.R-project.org/). Computations on genomic coordinate files and downstream analyses were conducted using the data.table R package v1.18.0 (data.table: Extension of data.frame; https://r-datatable.com).

After initial quality checks using FastQC, paired-end reads were trimmed using Trim Galore v0.6.10 (https://www.bioinformatics.babraham.ac.uk/projects/trim_galore/) and cutadapt v5.2 (MARTIN 2011) and aligned to the dm6 version of the *Drosophila* genome using Bowtie 2 v2.5.4 (LANGMEAD AND SALZBERG 2012) with the following parameter: --maxins 1000. Reads with a mapping quality <30 were removed using SAMtools v1.16.1 (LI *et al*. 2009). Peaks were then called for each replicate separately and from merged reads for each sample using MACS2 v2.7.7.1 (FENG *et al*. 2012) with the following parameters: -B --SPMR -g dm --keep-dup 1 -f BAMPE. For experiments on HA-tagged proteins, HA ChIP-seq and CUT&RUN data from the corresponding wild-type tissues were used as control files. For ChIP-seq experiments on endogenous proteins in 16–20 h embryos, input samples were prepared from the corresponding HA-tagged strains and used as controls. For CUT & RUN on endogenous PC protein in EDs, peak calling was performed without a control file. Only confident peaks called from merged reads and detected independently in both replicates were retained for further analyses. Confident peaks were defined as peaks with qValue > 5 and signalValue > 5. For each sample, MACS2 output pileup files were converted to bigWig coverage tracks for visualization.

### Clustering of ChIP-seq and CUT&RUN peaks

For each dataset, the union of confident peaks, as defined above, was merged and resized around peak centers to ±500 bp. A matched set of randomly sampled non-overlapping control regions was then generated. For each sample, mean coverage was computed for each region, log2-transformed after adding a pseudo count corresponding to the smallest non-zero value, and centered using the median value of the randomly sampled control regions.

The resulting normalized values across all peak regions were clustered using the supersom function from the kohonen R package v3.0.13 (WEHRENS AND KRUISSELBRINK 2018), using a 1 × 5 grid with topology = hexagonal and toroidal = TRUE. PcG categories were retrieved by overlapping peaks with the PcG binding-site labels defined in (LOUBIERE *et al*. 2020). For the quantification of Polycomb-mediated histone marks, ChIP-seq data were obtained from (LOUBIERE *et al*. 2020) (GSE126985) for H2AK118Ub in 14–16 h embryos and in EDs, (SCHUETTENGRUBER *et al*. 2014) (GSE60428) for H3K27me3 in 16–18 h embryos, and (LOUBIERE *et al*. 2016) (GSE36039) for H3K27me3 and SU(Z)12 in eye discs.

### Motif analysis of ChIP-seq and CUT&RUN clusters

JASPAR CORE Insecta motifs were counted within resized peak regions and randomly sampled control regions, as described above, using the motifmatchr R package v1.32.0 with the following parameters: bg = ‘genome’, p.cutoff = 1e-5, and genome = ‘dm6’. For each cluster, motif enrichment was assessed using a two-tailed Fisher’s exact test against the randomly sampled control regions, to determine whether the number of sequences containing at least one occurrence of a given motif was significantly higher in the test group. Multiple-testing correction was performed using the FDR method. Only the top five enriched motifs per cluster, with an FDR ≤ 1e-3 and a log2 odds ratio > 0 were displayed in the corresponding panels. All enriched motifs with FDR ≤ 5e-2 are provided in **Supplementary Table 2**.

### Immunoprecipitation (IP) of nuclear extracts for mass spectrometry analysis

*Drosophila* embryos from the PC-HA, SCE-HA and SU(Z)12-HA lines were dechorionated and extensively washed with EWB. Embryos were resuspended in hypotonic buffer (15 mM HEPES pH 7.6, 10 mM KCl, 5 mM MgCl₂, 0.1 mM EDTA, 0.5 mM EGTA, 350 mM sucrose, freshly supplemented with 2× protease inhibitor cocktail) using a volume equivalent to the embryo volume and homogenized in a pre-chilled Tenbroeck homogenizer (20 strokes). Lysates were centrifuged at 4,000 × g for 10 min at 4°C to separate cytosolic and nuclear fractions.

Nuclear pellets were flash-frozen until sufficient material had been collected. Nuclei were resuspended in 1.5 volumes of sucrose buffer (50 mM HEPES pH 7.0, 60 mM NaCl, 15 mM KCl, 10 mM MgCl₂, 0.34 M sucrose), and high-salt buffer (50 mM HEPES pH 7.0, 0.2 mM EDTA, 25% glycerol, 900 mM NaCl, 1 mM MgCl₂) was added dropwise to obtain a final NaCl concentration of 300 mM. Samples were incubated for 15 min at 4°C and next supplemented with 1 mM CaCl₂ and MNase (0.0125 U/µL; Sigma N3755) for digestion at 37°C for 15 min. Digestion was stopped by addition of EDTA to a final concentration of 25 mM, and extracts were incubated for 1 h at 4°C. The NaCl concentration was then adjusted to 150 mM, followed by ultracentrifugation at 49,000 rpm for 1 h at 4°C. The supernatant containing nuclear proteins was collected and protein concentration was determined using a BCA assay.

For immunoprecipitation, 1.5 mg of embryo nuclear extract was incubated overnight at 4°C with respectively 75 µL anti-HA high-affinity agarose (Roche, 11815016001). Embryo IPs were performed in a final volume of 1 ml adjusted with TEGN buffer (10 mM Tris-HCl pH 7.5, 0.5 mM EDTA, 150 mM NaCl, 10% glycerol). Beads were washed five times with wash buffer (10 mM Tris-HCl pH 7.5, 0.5% NP-40, 150 mM NaCl) and transferred to a new tube after the first wash. A final wash was done in 20mM of tris pH7.5 and 150 mM NaCl before samples were flash-frozen. Trypsin digestion was performed directly on the beads in 1M urea with 200 mM ammonium bicarbonate buffer (1 µg, 37°C, 8h, Promega cat # V5113). After digestion, peptide mix was acidified with formic acid and desalted with a MicroSpin C18 column (The Nest Group, Inc) prior to LC-MS/MS analysis. All experiments were carried out in biological triplicate. As a negative control, anti-HA IPs were performed using wild-type (WT) flies lacking an HA-tagged protein.

### Mass spectrometry (MS) analysis

Samples were analyzed using a LTQ-Orbitrap Fusion Lumos mass spectrometer (Thermo Fisher Scientific, San Jose, CA, USA) coupled to an EASY-nLC 1200 (Thermo Fisher Scientific (Proxeon), Odense, Denmark). Peptides were loaded directly onto the analytical column and were separated by reversed-phase chromatography using a 50-cm column with an inner diameter of 75 μm, packed with 2 μm C18 particles. Chromatographic gradients started at 95% buffer A and 5% buffer B with a flow rate of 300 nl/min and gradually increased to 25% buffer B in 52 min and then to 40% buffer B in 8 min. After each analysis, the column was washed for 10 min with 100% buffer B. Buffer A: 0.1% formic acid in water. Buffer B: 0.1% formic acid in 80% acetonitrile.

The mass spectrometer was operated in positive ionization mode with nanospray voltage set at 2.4 kV and source temperature at 305°C. The acquisition was performed in data-dependent acquisition (DDA) mode and full MS scans with 1 micro scans at resolution of 120,000 were used over a mass range of m/z 350-1400 with detection in the Orbitrap mass analyzer. Auto gain control (AGC) was set to ‘standard’ and injection time to ‘auto’. In each cycle of data-dependent acquisition analysis, following each survey scan, the most intense ions above a threshold ion count of 10000 were selected for fragmentation. The number of selected precursor ions for fragmentation was determined by the “Top Speed” acquisition algorithm and a dynamic exclusion of 60 seconds. Fragment ion spectra were produced via high-energy collision dissociation (HCD) at normalized collision energy of 28% and they were acquired in the ion trap mass analyzer. AGC and injection time were set to ‘Standard’ and ‘Dynamic’, respectively and isolation window of 1.4 m/z was used.

Information of the identified peptides and their corresponding proteins is provided in **Supplementary Table 3.**

### MS Data Analysis

Acquired spectra were analyzed using the Proteome Discoverer software suite (v2.3, Thermo Fisher Scientific) and the Mascot search engine (v2.6, Matrix Science). The data were searches against a Swiss-Prot UP_Drosophila (as in February 2020) plus a list of common contaminants and all the corresponding decoy entries. For peptide identification a precursor ion mass tolerance of 7 ppm was used for MS1 level, trypsin was chosen as enzyme and up to three missed cleavages were allowed. The fragment ion mass tolerance was set to 0.5 Da for MS2 spectra. Oxidation of methionine and N-terminal protein acetylation were used as variable modifications whereas carbamidomethylation on cysteines was set as a fixed modification. False discovery rate (FDR) in peptide identification was set to a maximum of 5%. Bona fide interaction proteins were obtained using the SAINT (Significance Analysis of INTeractome) software package with a FDR filter < 5%. The raw proteomics data have been deposited to the PRIDE repository with the dataset identifier PXD080680.

### Clustering of embryo MS hits

High-confidence MS hits were defined as hits satisfying the following criteria: log2 FoldChange > 3, BFDR ≤ 0.05, SaintScore > 0.8, TopoAvgP > 0.8, SpecSum ≥ 5, and NumReplicates ≥ 2. Among these, hits expected to be localized in the nucleus, defined as those with a Cellular Component annotation of ‘nucleus’, ‘chromosome’, were ranked based on their FoldChange values. Hits with a rank ≤ 12 in any of the three pull-downs were then clustered based on their log2 Fold Change values using k-means clustering in R with k = 6.

### Western blot (WB) experiments

Embryos were collected 16-20 h after egg laying. Following dechorionation, embryos were homogenized in radioimmunoprecipitation assay lysis (RIPA) buffer (50 mM Tris pH 7.5, 150 mM NaCl, 1% NP40, 0.5% Na-deoxycholate, 0.1% SDS, 2× protease inhibitor cocktail) using a Tenbroeck homogenizer and incubated on ice for 10 min. Samples were centrifuged at 10,000g for 10 min at 4 °C and the supernatant was transferred to a fresh tube. Protein concentration was quantified using a BCA protein assay and 10 µg of total protein was loaded per lane. Proteins were separated by SDS–PAGE for 40 min at 200 V in MES running buffer and transferred to membranes for 1 h at 1 A. Membranes were blocked for 1 h at RT in PBS containing 0.2% Tween and 10% milk powder at RT, incubated overnight at 4 °C with primary antibodies diluted in PBS containing 0.2% Tween-20 on a shaker, and subsequently washed in PBS containing 0.2% Tween-20. The following primary antibodies were used: rabbit anti-PC (1:2000) (SCHUETTENGRUBER *et al*. 2009), rabbit anti-SU(Z)12 (1:2000) (LOUBIERE *et al*. 2017), mouse anti-HA (HA (F-7), Santa Cruz catalogue no. sc-7392 and mouse anti-alpha-tubulin (1:5,000, DSHB, catalog no. AA12.1). HRP-conjugated secondary antibodies were incubated with the membranes for 2 h at RT. The following secondary antibodies were used: goat anti-rabbit HRP (1:15,000; Sigma-Aldrich, catalog no. A0545), rabbit anti-mouse HRP (1:15,000; Sigma-Aldrich, catalog no. A9044). Membranes were washed in PBS containing 0.2% Tween-20 and revealed using the Super Signal^TM^ West Dura Extended Duration Substrate kit (Thermo Fisher Scientific/Pierce). Chemiluminescent signals were acquired using a Bio-Rad ChemiDoc imaging system. WBs were quantified using Image Lab software v.6.1 (Bio-Rad).

### RT-qPCR experiments

For each genotype, 5 homozygous adult flies were collected in triplicates and homogenized in TRIzol reagent. Total RNA was extracted using RNA Clean & Concentrator kit (Zymo Research, catalog no. R1015) according to the manufacturer’s instructions and using an on-column DNAse I treatment (Qiagen #79254). 1 ug of total RNA was used for reverse transcription (RT) with the Maxima First Strand cDNA synthesis Kit for RT-qPCR (Thermo Fisher Scientific, catalog no. Ep0741). Quantitative PCR was performed on a LightCycler480 instrument (Roche) using the primers listed in **Supplementary Table 1.** Data were analyzed using the LightCycler software. Gene expression levels were normalized to the housekeeping gene *rp49*.

## Results and Discussion

### Construction of a complete set of HA-tagged core PRC1 components

To generate a panel of fly lines carrying endogenously tagged PcG proteins, we selected the HA epitope because it is a small 9 amino-acid tag that offers several advantages for multiple applications. In particular, HA is less sensitive to formaldehyde crosslinking than other commonly used epitopes, such as FLAG. Moreover, the availability of high-affinity nanoantibodies provides an excellent signal-to-noise ratio in antibody-based approaches. We chose to introduce the HA tag at the C-terminus of each protein, as our initial attempts to tag PSC at its N-terminal part produced viable but sterile flies (data not shown). Moreover, all core PRC1 components possess an invariable C-terminal region, allowing detection of all protein isoforms.

The CRISPR/Cas9 knock-in strategy and all primer sequences used in this study are summarized in **Fig. 1a** and **Supplementary Table 1**, respectively. gRNAs flanking the stop codon for each target gene were designed to generate a double-strand break and promote homology-directed repair (HDR). To facilitate endogenous tagging, we modified the previously published pHD-DsRed-attP donor vector (GRATZ *et al*. 2014) by inserting a cassette containing a 3xHA tag together with either 3xMYC or 3xFLAG tag upstream of the DsRed selection marker. Homology arms flanking the insertion site were cloned into the donor vector to enable C-terminal tagging of all core PRC1 members, including PC, PSC, SCE and both PH isoforms (PH-P and PH-D). In addition to the core PRC1 components, we tagged the PRC2 core subunit SU(Z)12 and several other chromatin-associated proteins acting either independently of PcG proteins, such as SU(HW) and CTCF, or antagonistically to PcG-mediated repression, such as Brahma (BRM) and Trithorax-related (TRR). A summary of all generated fly lines is shown in **Fig. 1b**.

**Figure 1:**
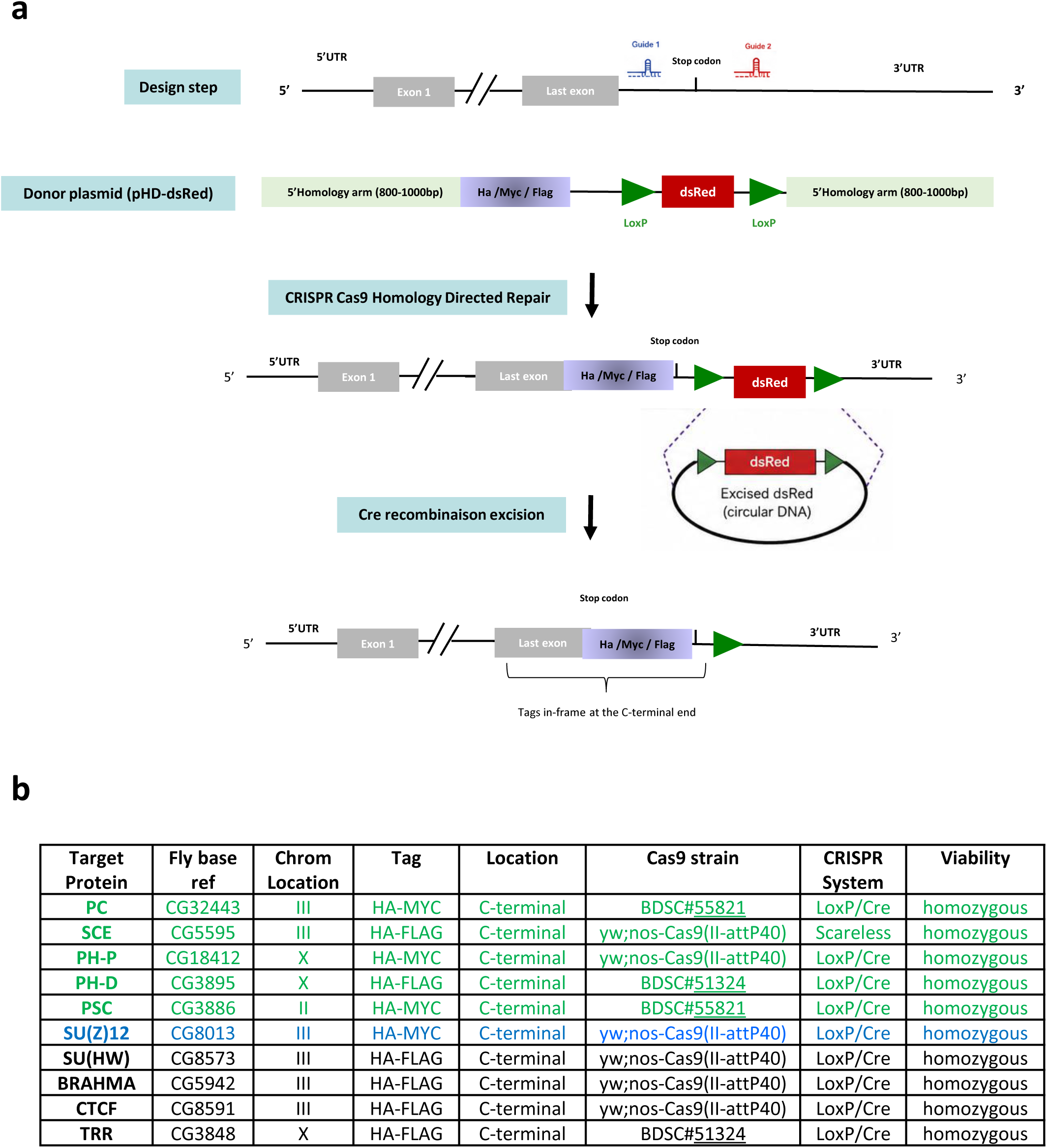
A complete HA-tagged toolkit for the core PRC1 complex and additional chromatin regulators. **(a)** Schematic representation of the strategy used to generate endogenous C-terminal HA-Myc (or HA-FLAG) fusion allele by CRISPR/Cas9-mediated genome engineering. Two gRNAs direct Cas9 cleavage immediately upstream and downstream of the endogenous STOP codon. Homology-directed repair (HDR) using a pHD-dsRED donor plasmid inserts an in-frame HA-Myc tag together with an LoxP-flanked dsRED selection cassette. Following identification of dsRED-positive knock-in alleles, Cre recombinase-mediated excision removes the dsRED cassette, leaving a clean C-terminal HA-Myc-tagged endogenous allele. The resulting edited allele preserves endogenous gene regulation while enabling detection of the native protein using anti-HA antibodies. **(b)** Table listing the generated tagged lines, including the target protein, FlyBase gene identifier, chromosomal location, epitope tag (HA-Myc or HA-FLAG), C-terminal insertion site, Cas9 background strain, CRISPR genome-editing strategy (Cre/LoxP or scarless), and viability of the final edited allele. All tagged lines were recovered as viable homozygotes.

After screening for DsRed-positive flies, the selection cassette was removed by crossing the flies to CRE-recombinase-expressing flies. Importantly, all resulting C-terminal tagged lines were homozygous viable and fertile. Correct insertion of the HA-tag was confirmed by PCR genotyping (**Supplementary Fig. 1**) and by sequencing of all edited loci.

Subsequent analyses focused on fly lines carrying tagged alleles of core PRC1 and PRC2 components. To determine whether endogenous tagging affected gene expression, we first quantified transcript levels by RT-qPCR (**Fig. 2a**). Importantly, no significant differences in mRNA levels were observed between tagged and untagged alleles for any of the PcG genes examined. Furthermore, Western blot analysis demonstrated that HA-tagging did not alter PC protein abundance (**Fig. 2b**). Together with the absence of detectable homeotic phenotypes in any of the tagged PcG lines, these results indicate that C-terminal HA-tagging does not significantly affect PcG gene expression or protein level.

**Figure 2:**
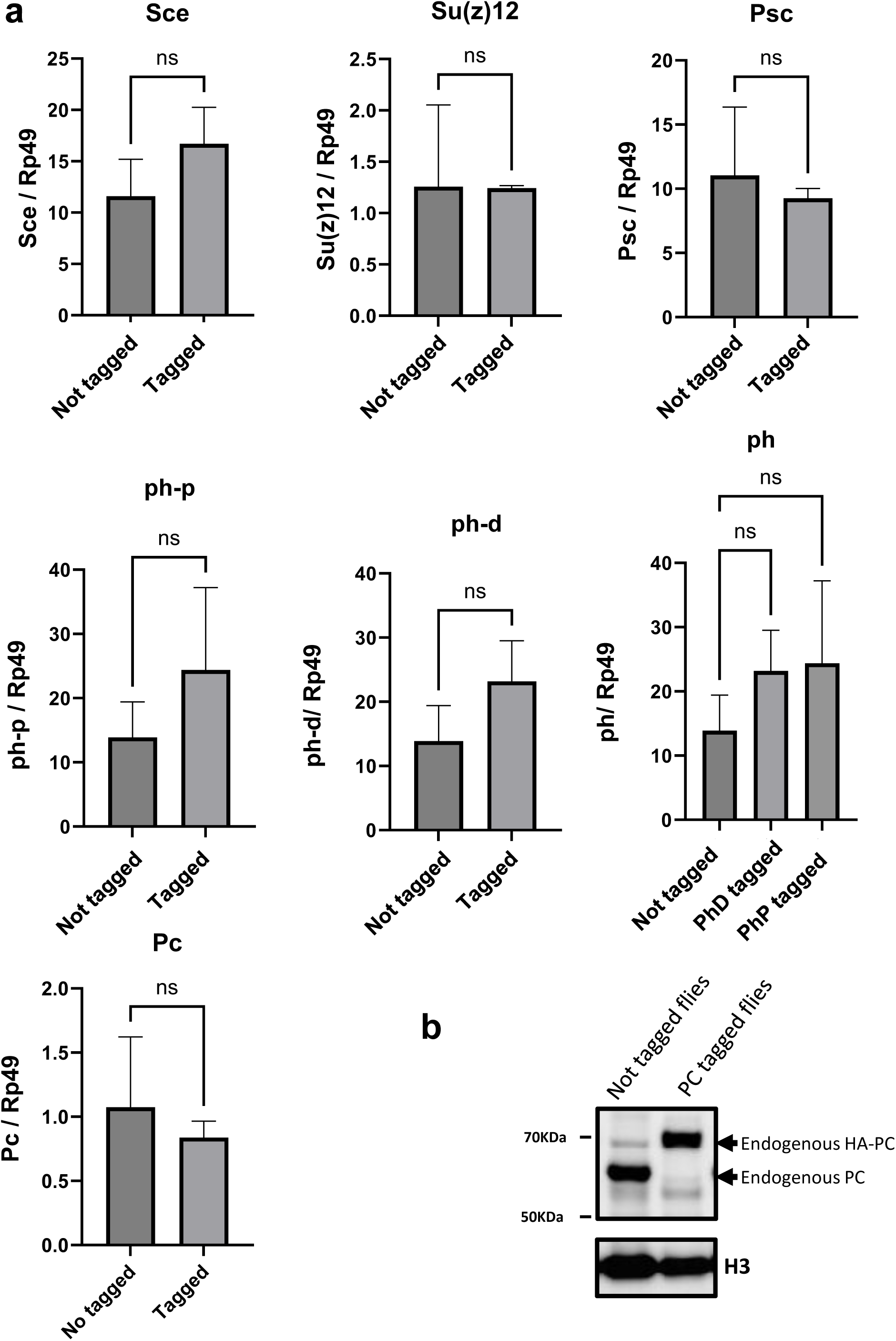
Endogenous HA-tagging does not alter PcG protein expression. **(a)** RT-qPCR analysis comparing transcript levels of the indicated endogenous untagged and HA-tagged alleles. Expression values were normalized to the housekeeping gene *Rp49*. Data represent three independent biological replicates. No significant differences in transcript expression were detected between untagged and HA-tagged lines (paired *t*-test; ns, not significant). **(b)** Western blot analysis of PC protein levels in untagged and HA-tagged fly lines. Comparable PC protein levels were observed in both genotypes, indicating that endogenous HA-tagging does not affect PC protein abundance.

### Endogenous HA tagging preserves chromatin targeting of PcG proteins

Having confirmed the correct expression of HA-tagged PcG proteins, we next investigated whether they are correctly recruited to their chromatin target sites. We first performed immunostaining experiments on polytene chromosomes from third-instar larval salivary glands using anti-HA and anti-PC antibodies. Previous studies have shown that all PRC1 and PRC2 core components, together with H3K27me3, colocalize at approximately 100 chromosomal bands (HAUENSCHILD *et al*. 2008). In agreement, PC staining in wild-type chromosomes revealed approximately 100 chromosomal binding sites, whereas anti-HA staining produced no detectable signal (**Fig. 3**, WT). Co-staining of chromosomes from HA-tagged PC fly lines with anti-HA and anti-PC antibodies revealed virtually perfect colocalization, indicating that the tagged protein is efficiently recognized by HA antibodies and localizes correctly to chromatin (**Fig. 3**, PC-HA line). Similarly, highly overlapping HA and PC staining patterns were observed in lines expressing HA-tagged PSC, PH-P, SCE, and SU(Z)12 proteins (**Fig. 3**), indicating that endogenous HA-tagging does not interfere with chromatin recruitment of PcG proteins. In contrast, the chromosomal distribution of HA-SU(HW) differed substantially from that of PC, confirming that SU(HW) binds to many genomic sites that are not bound by PC (**Fig. 3**). These observations further support the specificity of HA-tagged protein recruitment to their target chromatin and therefore the HA-tagging strategy.

**Figure 3.**
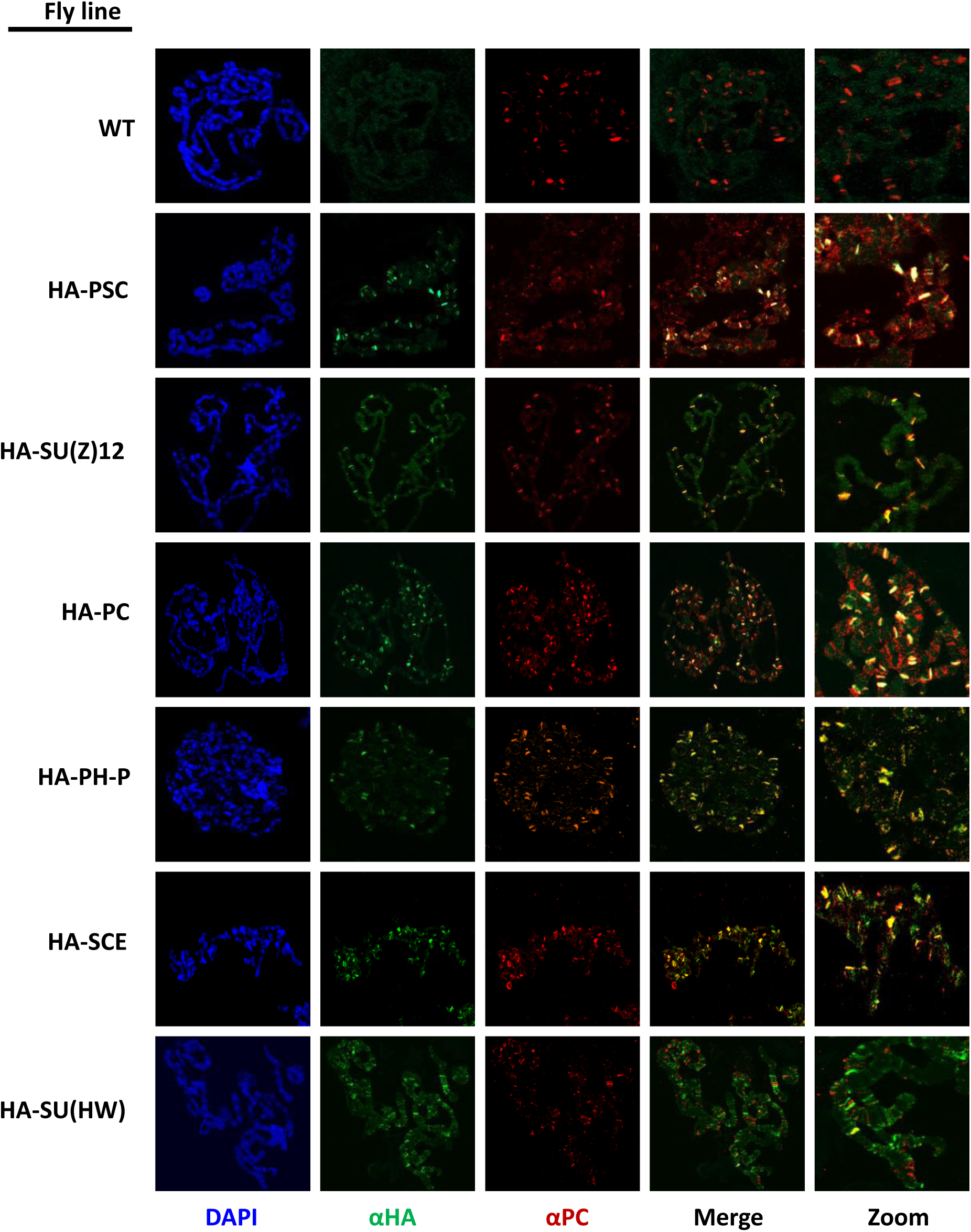
Chromatin localization of endogenously HA-tagged PcG proteins. Immunofluorescence staining of polytene chromosomes from third-instar larval salivary glands using anti-HA and anti-PC antibodies in the indicated fly lines. HA-tagged PcG proteins display chromosomal binding patterns that largely overlap with PC, indicating that endogenous HA-tagging preserves their normal recruitment to chromatin. In contrast, HA-SU(HW) exhibits only partial overlap with PC, consistent with its broader genomic distribution. No HA signal was detected in the wild-type control, which lacks an HA epitope.

To examine chromatin targeting of HA-tagged PcG proteins at a genome-wide scale, we next performed ChIP-seq experiments in *Drosophila* embryos. To this aim, we selected the HA-PC and HA-SU(Z)12 lines to compare the genome binding patterns of representative canonical PRC1 (PC) and PRC2 (SU(Z)12) components using the same HA antibody. Because well-characterized antibodies against the endogenous proteins are available, these experiments also enabled direct comparison of binding profiles obtained using anti-HA and endogenous antibodies. As a control, we additionally profiled HA-SU(HW), which is expected to bind distinct chromatin sites from those occupied by PcG proteins and to only partially overlap with them. As for PC and SU(Z)12, we compared SU(HW) binding profiles obtained with the endogenous SU(HW) antibodies and the HA antibodies. Finally, we analyzed HA-SCE, a component shared by canonical and non-canonical PRC1 complexes. The use of a common HA epitope allowed direct comparison of SCE chromatin occupancy with that of the canonical PRC1 subunit PC while eliminating potential biases caused by differences in antibody performance. This approach therefore provides an opportunity to assess the genomic distribution of PRC1 complexes containing SCE and to determine the extent to which cPRC1 and ncPRC1 occupy distinct or overlapping chromatin regions in *Drosophila* embryos (**Fig. 4a**).

**Figure 4.**
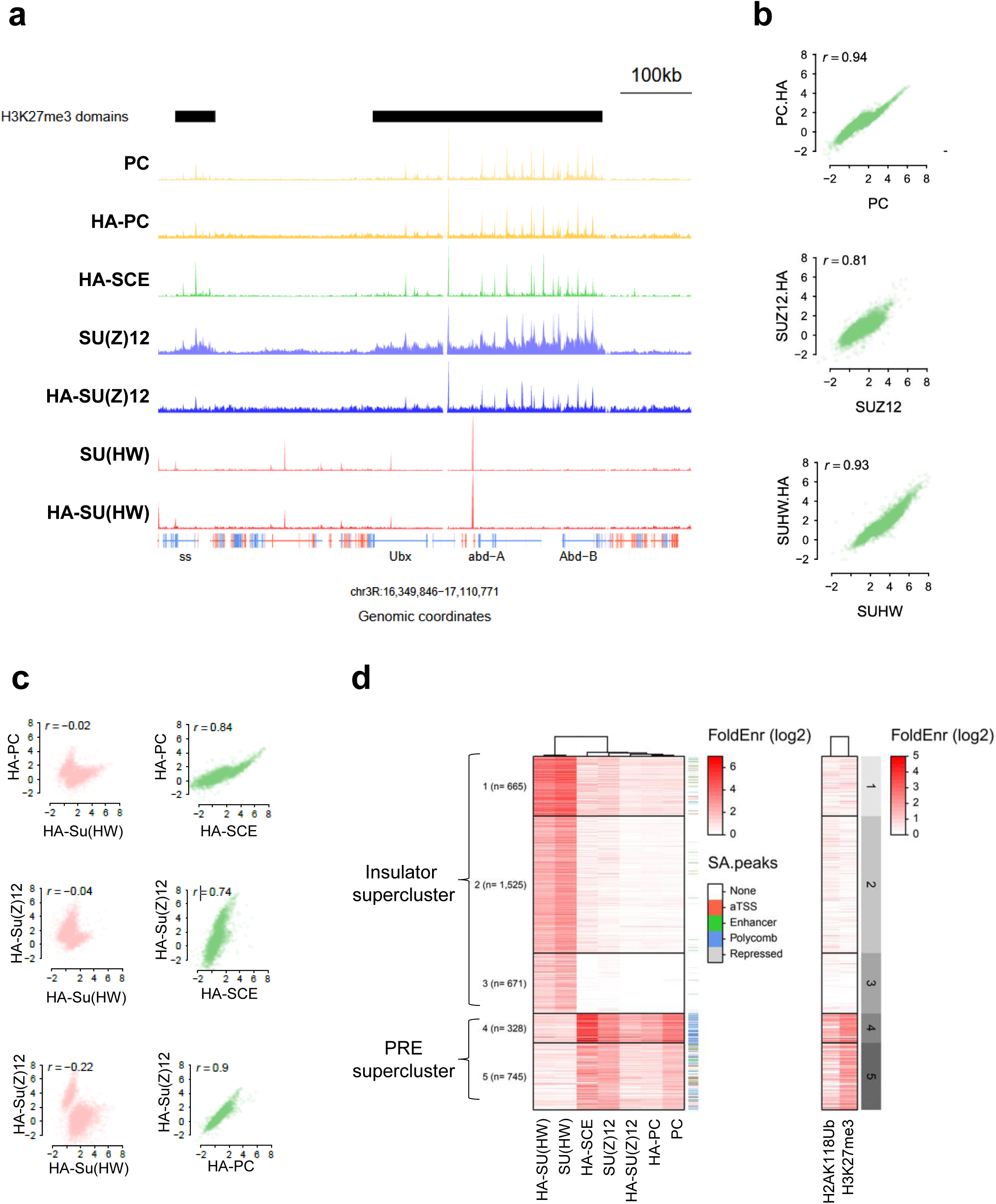
Genome-wide profiling of endogenously HA-tagged PcG proteins in *Drosophila* embryos. **(a)** Representative genome browser view showing ChIP-seq profiles of the indicated HA-tagged proteins across the bithorax complex (Bx-C) in *Drosophila* embryos. Chromatin immunoprecipitation was performed using either anti-HA antibodies or antibodies directed against the corresponding endogenous proteins. The highly similar enrichment patterns demonstrate that endogenous HA-tagging faithfully recapitulates the native chromatin-binding profiles of PcG proteins. **(b)** Scatterplots comparing ChIP-seq enrichment values obtained for the indicated HA-tagged proteins using either anti-HA antibodies or antibodies directed against the corresponding endogenous chromatin factors. **(c)** Scatterplots showing ChIP-seq enrichment values between the indicated HA-tagged proteins. **(d)** Clustering analysis of ChIP-seq peaks identifies distinct classes of genomic binding sites corresponding to insulators or PREs. The right panel shows the distribution of the histone modifications H2AK118ub and H3K27me3 across the same genomic regions in *Drosophila* embryos. Coloured annotations indicate the overlap of each cluster with previously annotated genomic features, including active transcription start sites (aTSSs), enhancers, Polycomb domains, and repressed chromatin states (LOUBIERE *et al*. 2020). The number of regions assigned to each cluster is indicated.

Genome-wide binding profiles obtained with anti-HA antibodies closely matched those obtained using antibodies directed against the corresponding endogenous proteins (**Fig. 4a**). Correlation analyses revealed strong agreement between the two datasets, with Pearson correlation coefficients of 0.94 for PC, 0.81 for SU(Z)12 and 0.93 for SU(HW) (**Fig. 4b**). Importantly, the binding profiles of HA-PC, HA-SCE and HA-SU(Z)12 were highly similar and strongly correlated with one another (**Fig. 4a,c**), confirming that most PRC1-bound regions are also occupied by PRC2 in *Drosophila* embryos (**Supplementary Fig. 2a**). By contrast, HA-SU(HW) binding showed little correlation with HA-PC binding (**Fig. 4a,c**), indicating that these two factors largely target distinct chromatin regions (**Supplementary Fig. 2a**).

Clustering analysis of ChIP-seq peaks identified a large “supercluster” corresponding to canonical PREs co-occupied by PRC1 and PRC2 (PC, SCE and SU(Z)12) (**Fig. 4d**). Notably, these regions displayed high enrichments for the H3K27me3 and H2AK118ub repressive histone marks and extensively overlapped previously annotated PREs and PcG binding sites in *Drosophila* embryos (LOUBIERE *et al*. 2020). Motif enrichment analysis further revealed over-representation of binding motifs for transcription factors known to contribute to PcG recruitment or PcG function, including GAGA factor (TRL), Combgap (CG), PHOL, and M1BP (SCHUETTENGRUBER AND CAVALLI 2009; ZOUAZ *et al*. 2017) (**Supplementary Figure 2b**). Interestingly, the PcG supercluster could be subdivided into two classes (Clusters 4 and 5) distinguished by their relative levels of PRC1 (SCE and PC) and PRC2 (SU(Z)12) occupancy. Cluster 4 exhibited higher levels of SCE and PC binding compared to cluster 5, suggesting differences in the relative contribution of PRC1 and PRC2 complexes to chromatin regulation at these sites. Finally, clustering analysis confirmed that most SU(HW)-bound regions were distinct from PcG target sites (**Fig. 4d**, CL1-3). These regions likely correspond to chromatin insulators. Consistent with this interpretation, they were strongly enriched for binding motifs recognized by Su(HW), CTCF, Pita, and CLAMP, all of which are known regulators of insulator activity (NEGRE *et al*. 2010; MAKSIMENKO *et al*. 2015; BAG *et al*. 2019) (**Supplementary Figure 2b**).

Next, we sought to test an alternative mapping approach to analyze the genome-wide binding profiles of HA-tagged proteins at a different developmental stage. To this end, we performed CUT&RUN (C&R) experiments in third instar larval eye-antennal imaginal discs (EDs) using anti-HA antibodies in HA-tagged PC, SCE, and SU(HW) lines. In parallel, we performed C&R experiments using antibodies against endogenous PC. As observed in *Drosophila* embryos, binding profiles obtained with anti-HA antibodies closely matched those obtained with antibodies directed against endogenous PC (**Fig. 5a**), with a Pearson correlation coefficient (r) of 0.83 between PC and HA-PC samples (**Fig. 5b**). Similarly, the binding profiles of HA-PC and HA-SCE were strongly correlated (r = 0.78; **Fig. 5a,b**), whereas SU(HW) binding profiles do not correlate with PcG-associated profiles (r = -0.48 and -0.34; **Fig. 5a,b**).

**Figure 5:**
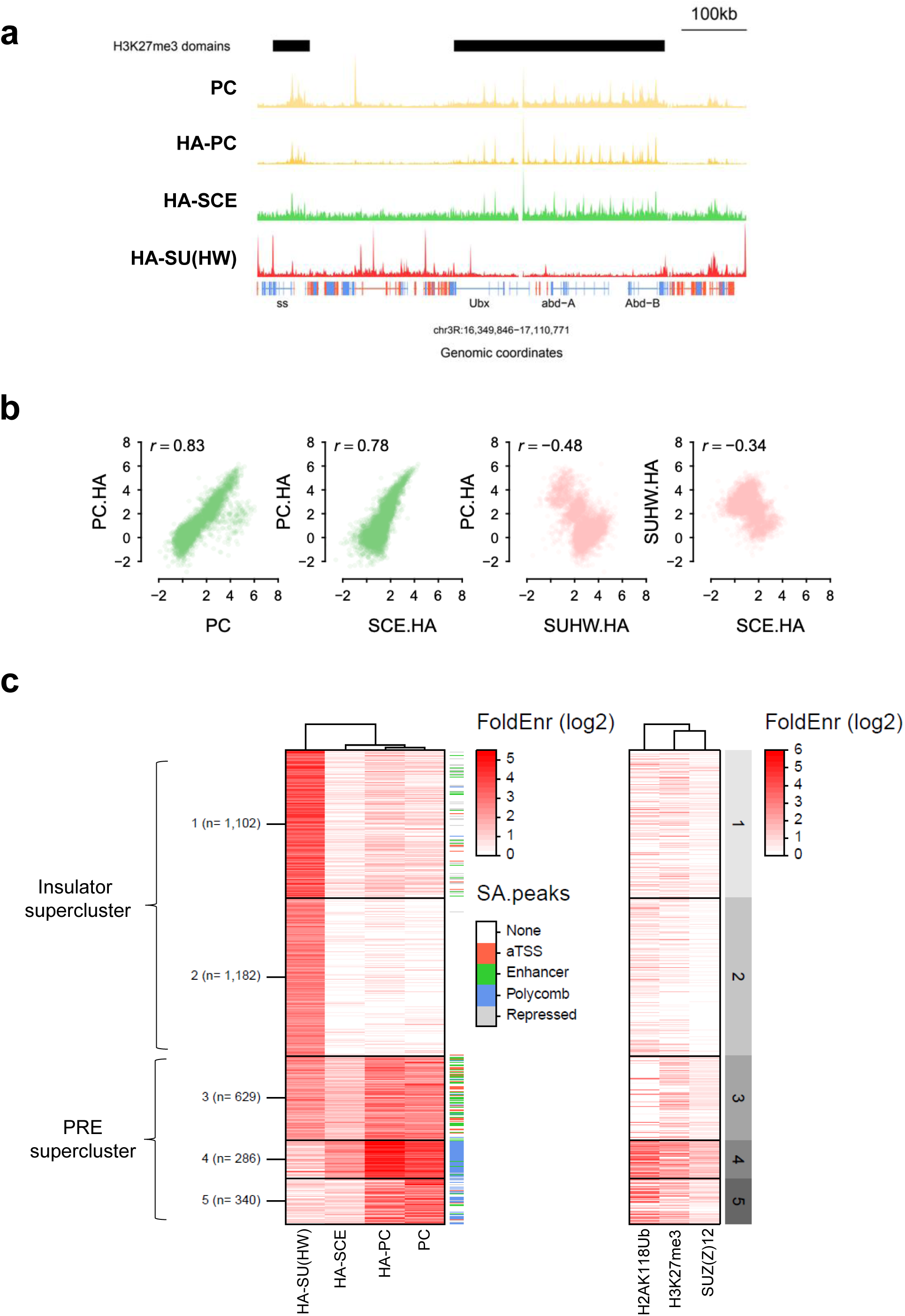
Genome-wide profiling of endogenously HA-tagged PcG proteins in *Drosophila* imaginal eye-antennal (ED) discs. **(a)** Representative genome browser view showing CUT&RUN profiles of the indicated HA-tagged proteins across the bithorax complex (Bx-C) in *Drosophila* EDs. CUT&RUN experiments were performed using either anti-HA antibodies or antibodies directed against the endogenous PC protein. **(b)** Scatterplots comparing CUT&RUN enrichment values for the indicated HA-tagged proteins obtained using either anti-HA antibodies or antibodies directed against the endogenous PC protein. The strong correlations between datasets confirm the reliability of HA-tagging for genome-wide chromatin profiling. **(c)** Clustering analysis of CUT&RUN peaks identifies genomic binding sites similar to those observed at the embryonic stage. The right panel shows the distribution of the histone modifications H2AK118ub, H3K27me3 and SU(Z)12 (LOUBIERE *et al*. 2020) across the same genomic regions in *Drosophila* EDs. Colored annotations indicate the overlap of each cluster with previously annotated genomic features, including active transcription start sites (aTSSs), enhancers, Polycomb domains, and repressed chromatin states (LOUBIERE *et al*. 2020). The number of regions within each cluster is indicated.

Importantly, clustering analysis of C&R peaks revealed the same “superclusters” previously identified in embryos (**Fig. 5c**). Clusters 1 and 2 correspond to SU(HW)-bound insulator regions and are enriched for binding motifs recognized by SU(HW) and other insulator-associated factors (**Supplementary Figure 2d**). By contrast, clusters 3–5 comprise PcG-bound genomic regions. Among these, clusters 4 and 5 correspond to canonical PREs and are characterized by high levels of the repressive histone marks H3K27me3 and H2AK118ub (**Fig. 5c**). As in embryos, these canonical PREs are associated with DNA binding motifs known to be implicated in PcG recruitment, including those recognized by (TRL, CG, Grainyhead (GRH) and PHOL (**Supplementary Figure 2d**). In contrast, cluster 3 lacks detectable H3K27me3 enrichment and corresponds to previously described neo-PRC1 target sites (LOUBIERE *et al*. 2016), which are associated with active promoters and enhancers specifically at this developmental stage. Consistent with this interpretation, comparison with previously published mapping data (LOUBIERE *et al*. 2020) showed that cluster 3 is largely devoid of SU(Z)12 binding (**Fig. 5c**), consistent with binding of PRC1 to regions that are not subjected to chromatin-mediated silencing. These genomic sites display a distinct DNA motif signature compared with canonical PREs, including increased enrichment for motifs recognized by the transcription factors M1BP, HR3, and Pannier (PNR). This observation suggests that these factors may contribute to the recruitment of PcG complexes to neo-PRC1 target sites in a developmental stage-specific manner (**Supplementary Figure 2d**). Interestingly, cluster 3 is also enriched for SU(HW) binding, supporting previous observations that SU(HW) localizes near a subset of PcG binding sites (NEGRE *et al*. 2010; SCHWARTZ *et al*. 2012) and can modulate PcG recruitment and PRE activity in transgenic contexts (COMET *et al*. 2011).

Notably, we did not identify a distinct cluster characterized by strong SCE enrichment in the absence of PC, nor did we detect a substantial number of SCE-bound regions lacking PC occupancy (**Supplementary Fig. 2a,c**), which would have been expected if ncPRC1 would bind to specific sites independent of cPRC1. This was true at both embryonic and larval stages. Together, these findings suggest that cPRC1 and ncPRC1 largely occupy a common set of PREs throughout *Drosophila* development. We note that these data are reminiscent of the substantial colocalization that was previously observed between cPRC1 and ncPRC1 at many genomic regions in mouse embryonic stem cells (MOREY *et al*. 2013), although a non-overlapping targeting at some specific loci was also reported in that study.

In summary, using complementary chromatin profiling approaches, we demonstrate that HA-tagged PcG proteins faithfully recapitulate endogenous chromatin binding patterns across different developmental stages.

### Identification of canonical and non-canonical protein interactors of HA-PcG proteins

Next, we used HA-tagged PC, SCE, and SU(Z)12 fly lines to identify protein partners of cPRC1, ncPRC1, and PRC2 components by HA-affinity purification followed by mass spectrometry (MS) in *Drosophila* embryos. The use of a common HA epitope for the purification of distinct PcG complexes enables cleaner and standardized comparisons between experiments. In addition, HA-based immunoprecipitation tolerates stringent washing conditions, thereby reducing background prior to MS analysis. Pull-downs were performed under native conditions (i.e., without crosslinking) on MNase-digested nuclear extracts to preferentially capture core complex components and direct interactors.

Immunoprecipitation of HA-tagged proteins from embryos recovered 36-40 peptides for HA-SU(Z)12, 35-39 peptides for HA-SCE, and only 5-8 peptides for HA-PC. Nevertheless, the pull-down enriched for PRC1 subunits at substantial levels. For subsequent analyses, we considered only interacting proteins annotated with Gene Ontology (GO) Cellular Component terms containing “nucleus” and “chromosome”. Applying this filtering strategy, we identified 113 high-confidence interactors for HA-PC, 136 for HA-SU(Z)12, and 13 for HA-SCE (see Methods for cutoff parameters) (**Fig. 6a, Supplementary Table 1**).

**Figure 6:**
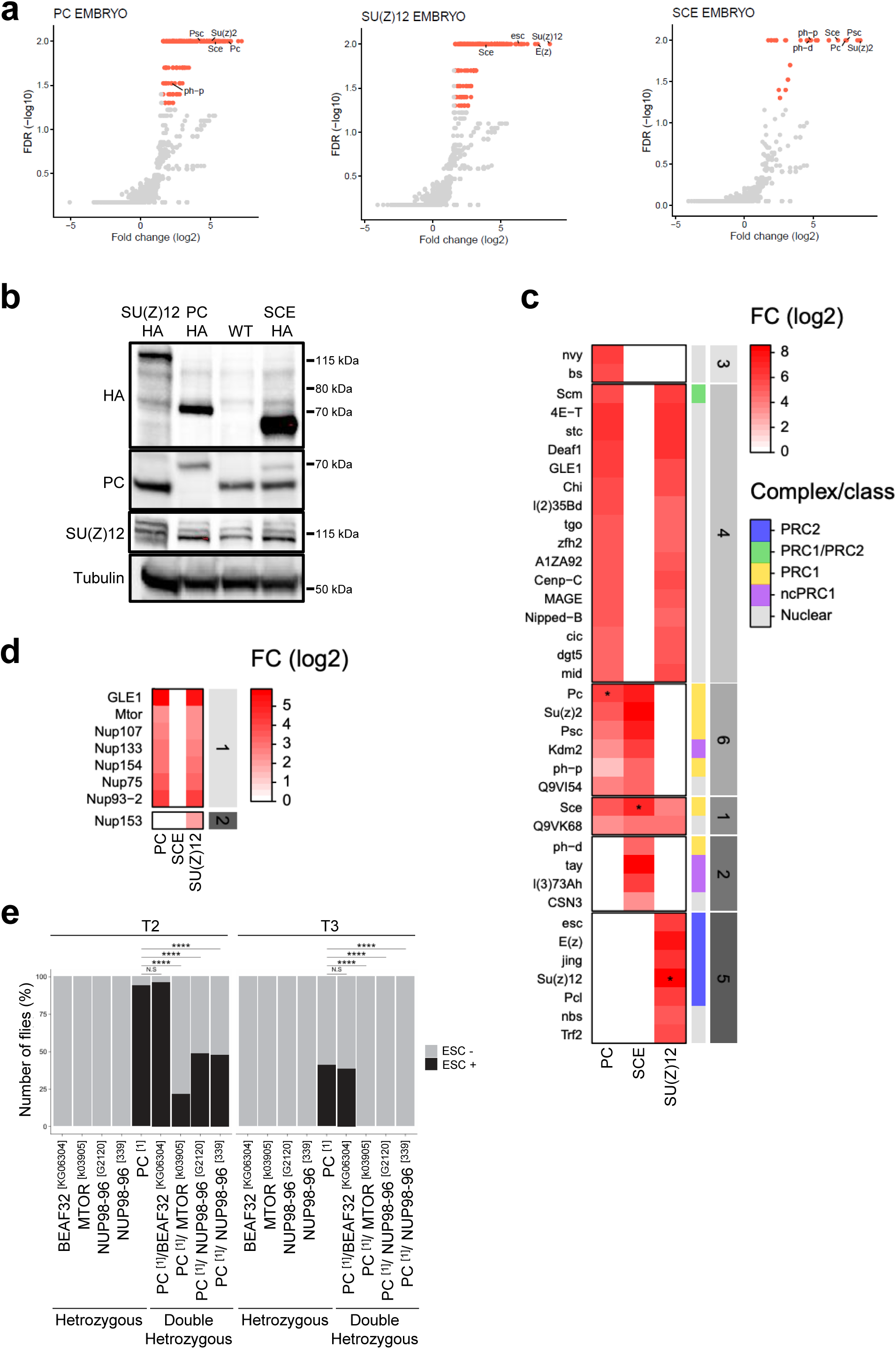
Identification of the native PcG protein interactome by HA immunoaffinity purification and mass spectrometry. **(a)** Scatter plots showing proteins significantly enriched in HA-PC, HA-SU(Z)12, and HA-SCE immunoprecipitations from *Drosophila* embryos relative to control samples. Significantly enriched interactors are highlighted in red. **(b)** Western blot analysis of HA-tagged protein expression levels in *Drosophila* embryos. Tubulin was used as a loading control. **(c)** Heatmap showing the top 12 nuclear hits identified for each HA-tagged protein. Hits were grouped using k-means clustering and clusters were ranked according to their center values. The annotation column on the right of the heatmap (complex/class) indicates known PcG complexes (KANG *et al*. 2022). **(d)** Heatmap showing enrichment of all annotated nuclear pore complex (NPC) proteins detected in the indicated HA-pull downs. **(e)** Genetic interaction analysis based on the homeotic Extra Sex Comb (ESC) mutant phenotype. The percentage of flies displaying the ESC phenotype is shown separately for the second (T2) and third (T3) pairs of legs. Statistical comparisons were performed against the control genotype *Pc*^1^ using Fisher’s exact test and P values were corrected for multiple testing using FDR: NS FDR ≥ 5x10−2, * FDR < 5x10−2, ** FDR < 1x10−2, *** FDR < 1x10−3, **** FDR < 1x10−5. Exact FDR values (in the order shown in the figure): 1x100, 3x10-12, 1x10-6, 8x10-7 (for T2) and 1x100, 4x10-6, 7x10-7, 5x10-7 (for T3). A minimum number of 25 flies were scored.

The comparatively low number of interactors identified with HA-SCE is unlikely to reflect reduced protein abundance in *Drosophila* embryos, as SCE is expressed at higher levels than either PC or SU(Z)12 at this developmental stage (**Fig. 6b**). Instead, this result may indicate that SCE engages in a more restricted set of stable interactions under native conditions, consistent with a more transient or catalytic mode of action within PRC1 complexes.

Clustering analysis of the top highest-confidence hits for each tagged protein using k-means clustering (see Methods) in embryos readily identified all known core canonical PRC1 components (PC, PH, SCE, and PSC) (**Fig. 6c**, yellow). In addition, several non-canonical PRC1 components (**Fig. 6c**, purple) were preferentially enriched in HA-SCE pull-downs compared to HA-PC pull-downs, despite the substantially fewer nuclear proteins detected overall in HA-SCE immunoprecipitations.

Previous BioTAP-XL studies in *Drosophila* embryos revealed the existence of distinct ncPRC1 complexes, including ncPRC1.1 and ncPRC1.3/5, characterized by the presence of KDM2 (ncPRC1.1) or TAY (ncPRC1.3/5), respectively (KANG *et al*. 2022). The identification of both KDM2 and TAY, together with L(3)73Ah, indicates that HA-SCE pull-downs co-purify multiple ncPRC1 complexes.

Similarly, all core PRC2 components (E(Z), ESC, and SU(Z)12) were specifically enriched in the HA-SU(Z)12 pull-down (**Fig. 6c**, blue). In addition, several PRC2 accessory proteins, including PCL and JING, were detected in association with SU(Z)12 but not in HA-PC or HA-SCE pull-downs. In contrast, SCM, which was previously shown to associate with both PRC1 and PRC2 (KANG *et al*. 2015), was specifically enriched in both HA-PC and HA-SU(Z)12 pull-downs (**Fig. 6c**, green).

Notably, despite applying stringent criteria to define bona fide interactors, we also identified several additional chromatin regulators as PRC1 and/or PRC2-associated proteins that have not previously been reported as core components of these complexes but are known to influence PcG function.

One such factor is Nipped-B, the *Drosophila* ortholog of NIPBL and a cohesin-loading factor. Cohesin has been previously shown to functionally interact with PcG proteins in the regulation of gene expression and to partially co-localize with PcG proteins on chromatin (SCHAAF *et al*. 2013; DORSETT 2019).

Another prominent interactor identified with both PRC1 and PRC2 is GLE1, a highly conserved mRNA export factor that functions at the nuclear pore complex (NPC). This observation prompted us to investigate whether additional NPC components were co-purified in our proteomic datasets. Indeed, we observed that multiple NPC proteins were significantly enriched in HA-PC and HA-SU(Z)12 pull-downs, including Megator (MTOR), NUP107, NUP133, NUP154, NUP75 and NUP93-2 (**Fig. 6d**). A functional connection between NPC components and PcG proteins has been reported previously. For example, NUP93 occupies strongly PcG bound loci and contributes to long-range chromatin interactions between Polycomb domains (GOZALO *et al*. 2020). In addition, several NPC proteins were identified in a genome-wide screen for factors affecting the nuclear distribution of PcG proteins (GONZALEZ *et al*. 2014).

To strengthen the functional link between PcG proteins and the nuclear pore complex (NPC), we tested genetic interactions between several NPC components and PcG proteins. Specifically, we established a genetic system in which PRC1-*Pc*^1^ mutant flies were crossed with mutants for either *Mtor* or *Nup98* to generate double heterozygous progenies (**Fig. 6e**). The *Pc*^1^ mutation causes homeotic transformations due to misregulation of the extra sex comb (*esc*) gene, resulting in the appearance of ectopic sex combs on the second and third pair of legs of male flies, whereas sex combs are normally restricted to the first pair of legs in wild-type animals. The penetrance of this phenotype depends on the extent of *esc* dysfunction. Consistent with this, mutation of the insulator protein BEAF-32 did not significantly affect phenotype penetrance (**Fig. 6e**). Strikingly, mutations in both *Mtor* and *Nup98* (the latter tested with two independent alleles) significantly reduced the penetrance of the extra sex comb phenotype (**Fig. 6e**). These findings provide genetic evidence for a functional interaction between PcG proteins and components of the NPC.

In summary, we generated a comprehensive collection of endogenously HA-tagged alleles for the core components of PRC1 together with additional chromatin regulators in *Drosophila melanogaster*. We demonstrate that these alleles faithfully recapitulate endogenous protein expression and chromatin binding, providing a standardized platform for chromatin profiling and proteomic analyses across developmental stages. Application of this toolkit revealed that canonical and non-canonical PRC1 complexes largely occupy the same genomic targets during development. In addition, proteomic analyses recovered known PcG complex components and identified previously unrecognized PcG-associated factors, including multiple components of the nuclear pore complex. Combined with genetic interaction studies, these findings support a functional connection between Polycomb-mediated gene regulation and the nuclear pore complex.

Together, our work establishes a versatile experimental framework for dissecting the composition, chromatin occupancy, and regulatory functions of Polycomb complexes and other chromatin-associated protein networks in vivo. Beyond the biological insights reported here, this resource provides a standardized platform for reproducible and integrative studies of PcG function and should facilitate the systematic analysis of chromatin regulatory complexes by the broader research community.

## Data Availability

All genomic data have been deposited on GSE338868, accessible at https://www.ncbi.nlm.nih.gov/geo/query/acc.cgi?acc=GSE338868 (reviewer token yrspuegezvuxtsb) and is publicly available as of the date of publication. Any additional information required to reanalyze the data reported in this paper is available from the corresponding author upon request. All proteomics data have been deposited in the PRIDE repository under identifier PXD080680, accessible at https://www.ebi.ac.uk/pride/login (reviewer token lzjP9QzSDWQe). All fly lines are available on request.

## Acknowledgments

We are grateful to Elissa P. Lei for the generous gift of anti-SU(HW) antibodies. We would like to thank Montpellier Ressources Imagerie (MRI), the PPM-FPP proteomics platform for advice and project discussions and the *Drosophila* facilities (BioCampus Montpellier). We also acknowledge the support of CNRS, INSERM, and of the University de Montpellier.

## Author Contributions

G.C., L.F., A-M.M. and B.S. conceived this study. L.F. generated mutant fly lines. L.F., A.D., A-M.M. and B.S. performed experiments. V.L. performed bioinformatic analysis of ChIP-seq, MS and CUT&RUN experiments. J.M-S. and E.S performed MS experiments and bioinformatic analysis of MS experiments. B.S. and L.F wrote the manuscript with the input of all other authors.

---

**Supplementary Figure 1.**
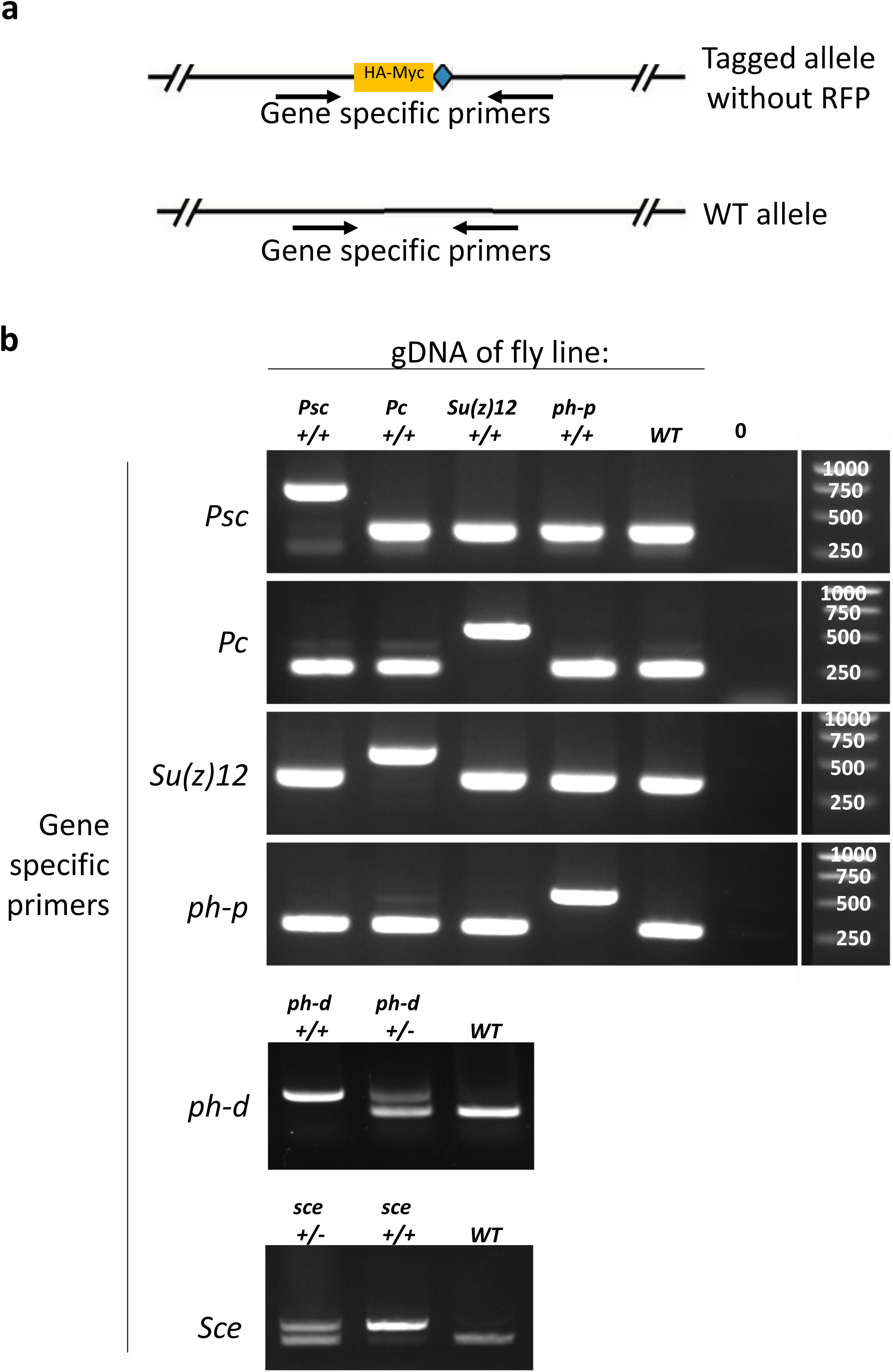
Genotyping of endogenously HA-tagged PcG alleles. **(a)** Schematic representation of the genotyping strategy, indicating the positions of the PCR primers used to distinguish wild-type and HA-tagged alleles. Insertion of the HA tag increases the size of the PCR product relative to the corresponding wild-type allele. **(b)** Representative agarose gel electrophoresis of the indicated fly lines, confirming successful insertion of the endogenous HA tag. The observed band sizes are consistent with the expected PCR products for the HA-tagged alleles.

**Supplementary Figure 2:**
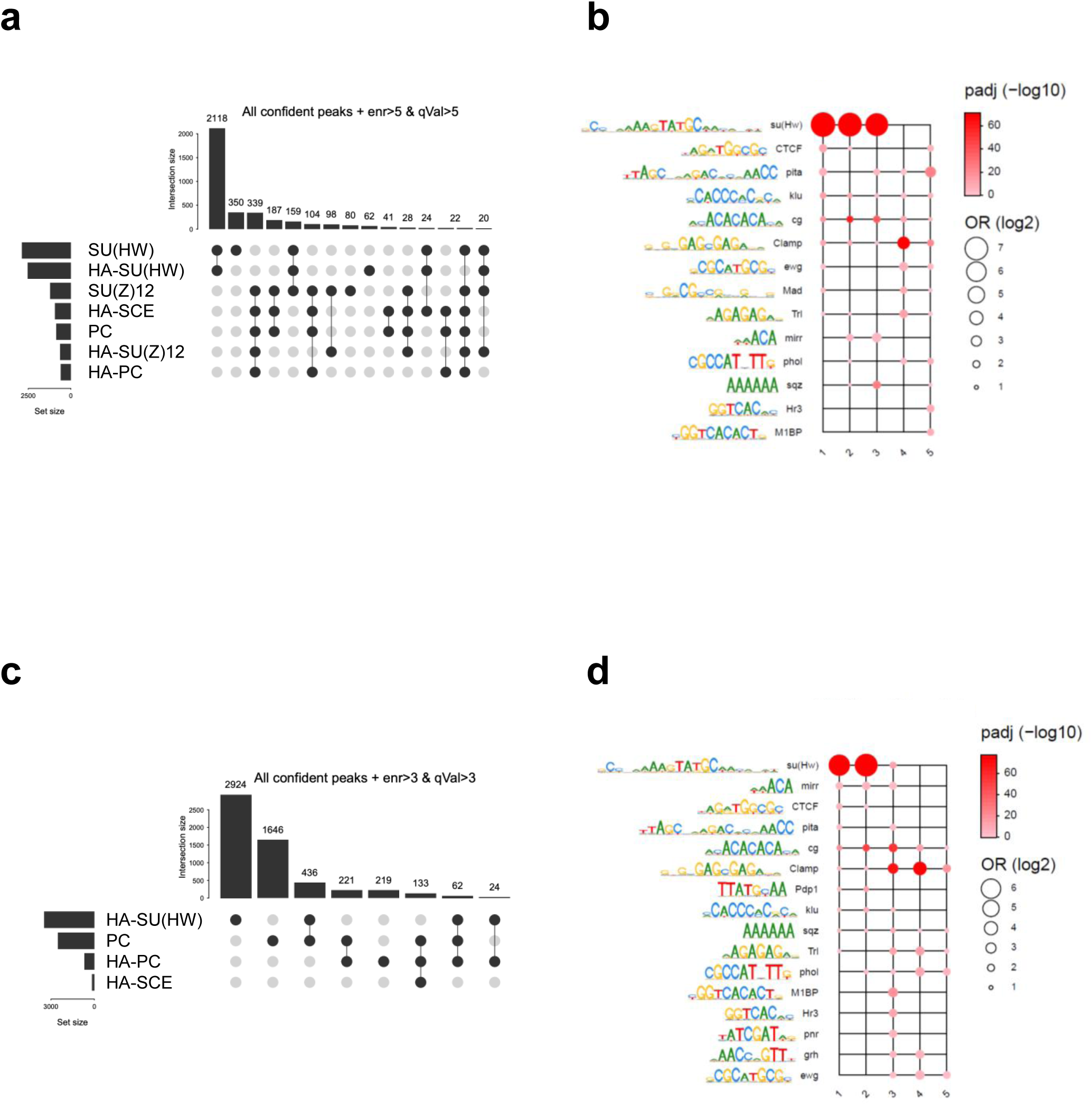
Overlap of genomic binding sites among endogenously HA-tagged PcG proteins and DNA motif analysis. **(a)** UpSet plot showing the intersections between ChIP-seq peak sets for the indicated HA-tagged proteins in *Drosophila* embryos. Horizontal bars indicate the total number of peaks identified for each factor, whereas vertical bars represent the number of genomic regions shared among the indicated combinations of factors. **(b)** Enrichment analysis of transcription factor binding motifs within the five genomic clusters identified by ChIP-seq. Representative sequence logos for enriched motifs are shown on the left. Circle color indicates the statistical significance of motif enrichment (adjusted *P* value, −log10 scale), whereas circle size reflects the magnitude of enrichment (log2 odds ratio) **(c)** UpSet plot showing the intersections between CUT&RUN peak sets of the indicated HA-tagged proteins in *Drosophila* eye-antennal imaginal discs. **(d)** Enrichment analysis of transcription factor binding motifs within the five genomic clusters identified by CUT&RUN.

**Supplementary Table 1:** List of primers, antibodies and *Drosophila* lines used or created in this study.

**Supplementary Table 2:** Enriched DNA motifs identified in ChIP-seq and CUT&RUN data sets

**Supplementary Table 3:** Summary of peptides and proteins identified in MS analysis

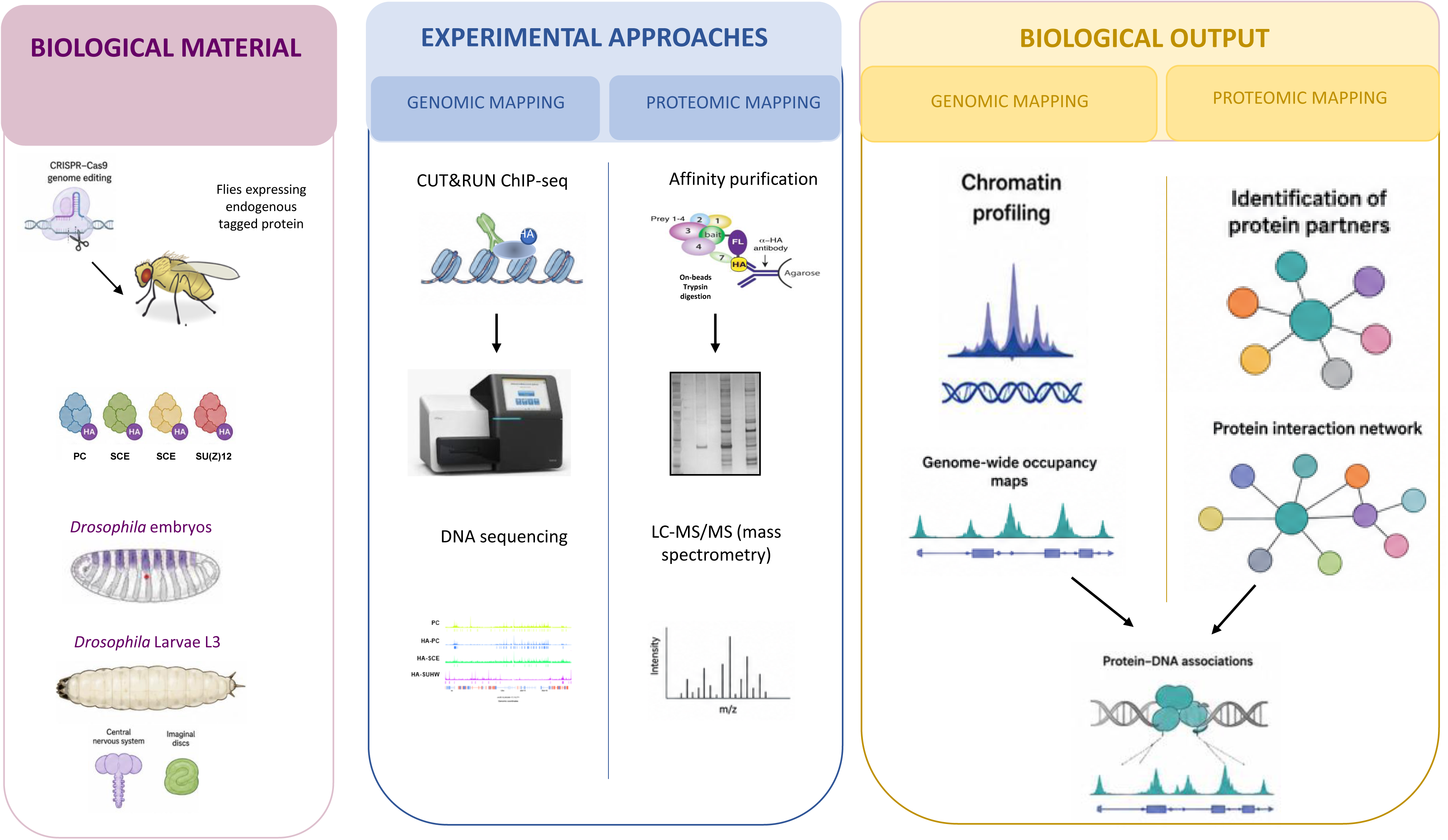

